# Steering in the presence of a gaze-contingent occlusion over a quarter of the visual field

**DOI:** 10.1101/2025.03.02.641038

**Authors:** Arianna P. Giguere, Matthew R. Cavanaugh, Krystel R. Huxlin, Duje Tadin, Brett R. Fajen, Gabriel J. Diaz

## Abstract

Why do some cortically blind (CB) drivers who are missing vision from a quadrant or hemifield have trouble maintaining a central lane position, while others do not? A recent driving study in virtual reality showed that most patients with right-sided visual field deficits (right CB) perform similarly to controls, while most of those with left CB demonstrated a unique pattern of steering biases (Giguere et al., 2025). In this study, we tested the hypothesis that these biases could result from loss of visual information falling on the blind field. The steering and gaze behavior of 24 subjects with normal vision (mean age: 19.8 years, SD: 1.44) were recorded in a virtual reality steering task while gaze-contingent occluding masks were imposed on a quadrant of their visual field. The central five degrees of vision were spared to mimic the sparing present in most CB patients. Turn direction (left/right), turn radius (two non-constant radii), and occlusion quadrant (one of four quadrants or no occlusion) were randomized between trials. We found that the pattern of steering biases observed in CB drivers were not replicated when visually-healthy drivers were subjected to gaze-contingent masks, and we conclude that it may be a mistake to characterize the effects of cortical blindness on steering behavior as consistent with a simple omission of visual information. This insight has the potential to guide future research on CB adaptation to their visual impairments and possible interventions to improve their steering performance.

## Introduction

Vision is fundamental in the guidance of steering to maintain lane position when driving. Although it is unsurprising that stroke-affected individuals with a loss of conscious vision in one quarter to one half of their visual field exhibit biases in lane position (Bowers, Mandel, Goldstein, & Peli, 2010) when steering, it is notable that these biases do not affect *all* individuals with cortical blindness (CB) equally; a recent study showed that the lane positioning of some CB patients in response to changes in visual motion information is nearly unaffected relative to controls, while it is greatly affected in others (Giguere et al., 2025). Why do some CB drivers have trouble maintaining lane position while others do not? An understanding of how CB affects visuo-motor tasks like driving is crucial knowledge that can be used to inform future vision rehabilitation methods.

Emerging evidence from controlled studies of driving behavior paints an unclear picture of whether the location of the blind field affects steering behavior. A rigorous on-road driving evaluation of 10 CB patients revealed that of the four patients who failed the assessment, all of them had a left hemifield deficit (Kasneci et al., 2014). Similarly, in a recent study involving a steering task in virtual reality, CB participants with a blind area on the left side of their visual field (left CB) demonstrated different lane position biases than those with right CB (Giguere et al., 2025). Analysis indicated that left CB drivers adopt lane positions that are biased away from their blind field when navigating turns, but those with CB on their right side do not (Giguere et al., 2025). ^1^ In the same study, a within-subject manipulation of optic flow revealed another asymmetry in behavior between left and right CB: the left CB group was relatively insensitive to changes in optic flow density compared to those with a blind area on the right side of their visual field (right CB) and visually-healthy participants (Giguere et al., 2025). Giguere et al. speculated that the relative insensitivity of left CB to optic flow was reflective of a disruption to motion processing caused by the stroke damage to the *right brain hemisphere*. This hypothesis is aligned with independent results that suggest the right brain hemisphere may play a dominant role in the processing of motion information (Alipour & Kazemi, 2015; Bosworth & Dobkins, 1999; Barthélémy & Boulinguez, 2002).

In contrast, a few studies have failed to find an effect of visual deficit location on steering. In a driving simulator where participants with CB could control their own speed, Szlyk, Brigell, and Seiple (1993) found no correlation between visual field loss location and lane position, though they did note that all patients had increased lane variability. Similarly, Bowers et al. (2010) reported that CB drivers were biased in lane position away from their blind fields in a driving simulation with oncoming traffic and off-road pedestrians, regardless of whether they had left or right CB. Their task introduced complex driving elements with the potential to obscure any between-group differences. Additionally, it is also possible that their sample size of six left CB and six right CB participants was too small to detect significant between-group differences, even if differences existed. In a more recent study, Biebl, Arcidiacono, Kacianka, Rieger, and Bengler (2022) developed a virtual driving simulation featuring oncoming traffic and a gaze-contingent simulation of CB (a full hemisphere of occlusion representing either left CB or right CB). They found that all participants showed reduced lane position stability and greater absence of large gaze movements under these simulated visual field loss conditions. There were no significant differences in average lane position between the left CB and right CB groups, although their simulated CB occlusions were limited in their generalizability to real CB, as they lacked foveal sparing, which is present in most CB patients (Horton, Economides, & Adams, 2021) and dramatically impacts both gaze patterns and navigation (Wood, Black, Mallon, Kwan, & Owsley, 2018; Patterson, Howard, Hepworth, & Rowe, 2019).

The present study was designed to address the uncertainty from previous work regarding the specific effect of CB deficit location on driving performance. Specifically, participants were immersed in a virtual reality environment and tasked with steering to maintain a central lane position within the winding single-lane roadway while *gaze-contingent* occlusions were placed over one quadrant of the visual field at a time, in a manner that spared foveal vision. Our goal was to test whether this manipulation of visual information in visually-healthy individuals could replicate the asymmetric steering biases observed previously (Giguere et al., 2025; Kasneci et al., 2014) in left vs. right CB patients. The present study is the first of its kind to test the impact of losing gaze-contingent visual information from each of the four visual field quadrants separately while steering in VR. Leveraging gaze-contingent methods was essential to promote ecological relevance as it pertains to CB patients, whose visual impairments are naturally fixed in retinal coordinates and follow the eyes. This method also allowed participants to make free head and eye movements, allowing us to analyze potentially compensatory gaze strategies (Bowers, 2016; Parker et al., 2011; Bahnemann et al., 2015; Biebl et al., 2024; Iorizzo, Riley, Hayhoe, & Huxlin, 2011; Bowers, Ananyev, Mandel, Goldstein, & Peli, 2014; Papageorgiou, Hardiess, Mallot, & Schiefer, 2012; Biebl & Bengler, 2021) that might be exhibited in the presence of a simulated occlusion mimicking CB.

When forming predictions about the effects of a gaze-contingent mask on steering behavior, it is helpful to first characterize some pre-existing steering biases in visually-healthy drivers. Even when instructed to maintain a central lane position, such drivers in a prior study were biased to position themselves towards the left road edge (Giguere, Huxlin, Tadin, Fajen, & Diaz, 2024). This behavior is consistent with the position of the steering wheel and driver on the left side of U.S. vehicles which are driven on the right side of the road. The finding mirrors that of Learmonth, Märker, McBride, Pellinen, and Harvey (2018) who found that people used to driving on the left side of the road are biased in the opposite direction. To account for these biases within the lane and to pinpoint the effect of the mask on steering despite these pre-existing tendencies, our predictions concern the steering behavior of each participant relative to their own steering behavior without a mask. We will hereafter refer to this form of bias as control-relative bias, or *bias*_*cr*_. ^2^

Independently of the aforementioned steering biases shared by all drivers, the bias_*cr*_ demonstrated by those with left CB has been shown to be asymmetric with respect to the bias_*cr*_ of right CB (Giguere et al., 2025). To remind the reader, Giguere et al. (2025) found that left CB participants were biased away from their blind fields relative to age-matched controls, but right CB participants were not. A finding that visually-healthy participants exposed to corresponding gaze-contingent masks demonstrate a similar steering asymmetry would support the interpretation that the asymmetry can be attributed to the omission of visual information rather than a result of differences in the way that information might be processed following cortical damage. Alternatively, if the bias_*cr*_ of visually-healthy individuals is symmetric in the presence of left and right visual masks, this would suggest that a lack of visual information alone cannot explain the steering asymmetry observed in CB patients.

Considering the evidence summarized thus far, we predict that visually-healthy participants with artificially imposed gaze-contingent blind fields will be equally biased in lane position away from occlusions on the left and right sides of their visual fields, unlike participants with cortical damage who completed a similar task (Giguere et al., 2025). Contrary to our prediction, if steering in the presence of a left visual occlusion causes lane position biases away from the simulated blind fields but steering with a right visual occlusion does not, our results would support the interpretation that the steering asymmetries found in previous studies (Giguere et al., 2025; Kasneci et al., 2014) are more likely to be caused by the visual occlusion of information rather than the influence of cortical damage on processes used for navigation. The findings will provide new insights into how quadrant-specific visual occlusion affects steering, and they may also guide future research on adaptation and intervention strategies for CB individuals.

## Methods

### Participants

Thirty participants between the ages of 18 and 24 were recruited for this study (mean age: 19.8 yrs, standard deviation 1.44, 18 self-reported as male, 11 as female, and one as nonbinary). All participants had normal or corrected to normal vision with contacts. Glasses were not allowed because they did not fit inside the head-mounted display. Only five participants wore contacts, and all except three participants possessed a driver’s license. The majority of participants reported that they drove at least once a week, while the remaining 9 drove occasionally (7) or never (3). Three subjects were unable to complete the task due to experiencing VR sickness, and they were excluded from analysis. One of the participants without a license had notably different steering results than others, and two more with licenses had a difficult time following instructions and/or blinked at a rate that led to poor eye tracking quality. These three were removed from analysis as well, leaving a total of 24 participants.

In the results section, we report the data from a follow-up study in which the mask location was fixed across blocks instead of being presented in a randomized location on each trial. These participants were subject to similar recruiting guidelines. This experiment involved eight participants, (mean age: 20.0 yrs, standard deviation 1.11, 5 self-reported as male, 3 as female).

Both studies were approved by RIT’s institutional review board ethics committee. Informed consent was obtained from all participants, and all data collection was performed in accordance with the guidelines and regulations defined in the consent forms. Participants were compensated in class credit for their time except for those participating in the follow-up study who were paid $10 instead.

### Apparatus

The virtual environment was developed on a computer with an AMD Ryzen 9 5950X 16-core processor, an NVIDIA GeForce RTX 3080 Ti graphics card, and 32.0 gigabytes of RAM. We utilized a Vive Pro Eye headset to display the environment which includes an integrated eye tracking system. The Vive Pro Eye has a 110° FOV, and frame rate 90 Hz (VIVE, n.d.), thus the gaze data and all other time-dependent metrics were sampled at this same frequency. The 3D experiment environment was developed in Unity3D (v2021.3.0f1), with SRanipal (v1.3.6.12) and SteamVR with two Lighthouse Base stations (v1). The OpenVRLoader and OpenXR options were selected under Unity’s XR plug-in management settings. The block and trial structure of the environment was made possible using the Unity Experiment Framework (UXF (Brookes, Warburton, Alghadier, Mon-Williams, & Mushtaq, 2020)).

A Logitech G920 steering wheel (Logitech, 2025) was used to control steering in the virtual environment. The Logitech software allows a user to change wheel sensitivity and centering spring strength, and we chose a sensitivity of 40 and spring strength of 40 with the goal of promoting both comfort and control for participants. The sensitivity setting adjusted the calibration between wheel angle and angle turned within the virtual environment, while the spring strength changed the strength of the wheel’s resistance back to its orientation at rest.

Data analysis occurred in an Anaconda virtual environment of Python version 3.9 and utilized packages including Numpy (v1.26.4), Pandas (v2.2.2) for data analysis and Matplotlib (v3.8.4) and Seaborn (v0.13.2) for data visualization.

### Task environment

Participants were immersed in a three-dimensional virtual environment where they were tasked with using the steering wheel to maintain a central position within a one-lane continuous roadway while traveling at a constant speed (19m/s). The ground plane was heavily textured with a 2D textured-grass pattern. The path was defined by a dark shading in high contrast to the surrounding grass pattern (see Figure 1). This decision, as well as the decision to have no visible virtual car dashboard or wheel was deliberately made to increase the strength of the visual motion information from the ground plane upon movement. Trees were randomly positioned on both sides of the road at a constant density across all trials and appeared prior to each turn, with the effect that they added additional motion information to the upper visual field.

**Figure 1:**
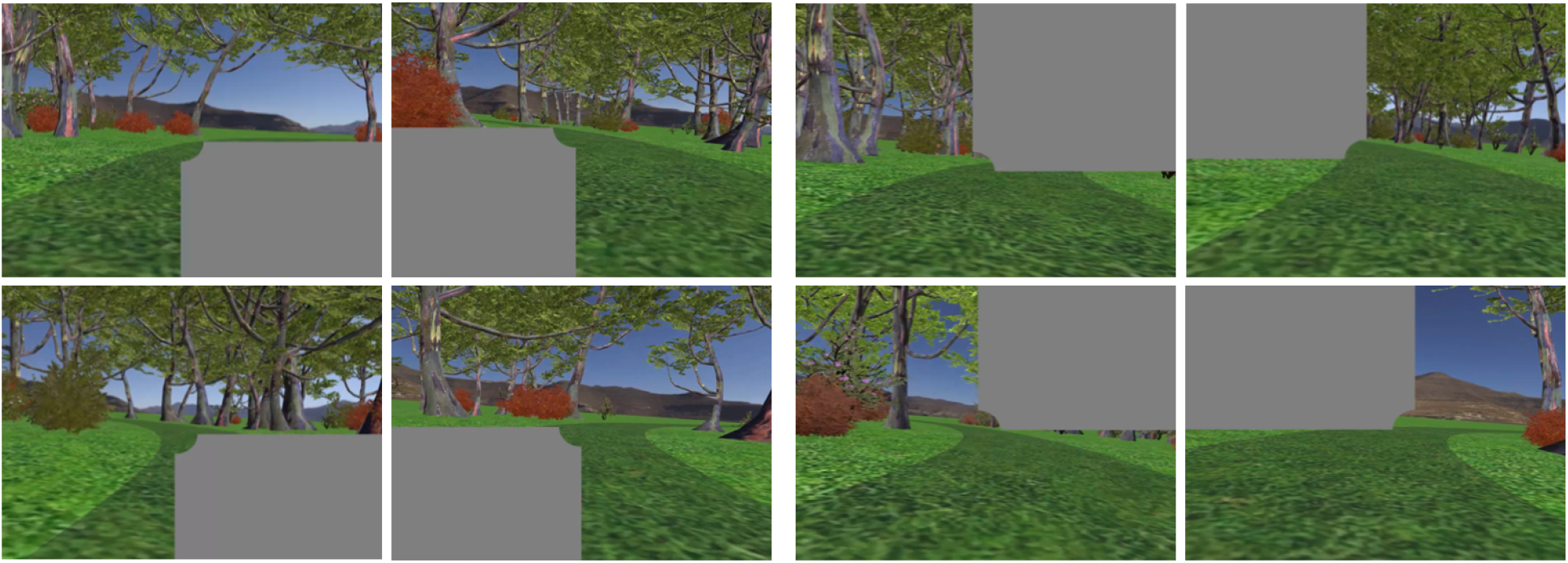
Eight images of the task environment as seen from inside the headset. The field of view is cropped significantly for viewing purposes. The gray occlusion was placed according to where the participant was looking at each moment in time. Vision in the foveal area is spared with a radius of 2.5°, as seen by the small quarter-circle cutout in each mask. The left two columns are examples of turns towards and away from a simulated deficit in the lower visual field, while the right two columns display examples of turns towards and away from a simulated deficit in the upper visual field. Not shown are the control trials where no mask is present. While the exact gaze locations of individual participants varied, these screenshots generally illustrate the amount of the road visible when participants navigated the curves.

#### Road generation

The road was constructed on-the-fly so that it always extended two turns ahead of the turn currently occupied by the participant, and each one was removed after it had been traversed. This process was done seamlessly and provided the sensation of a never-ending continuous roadway. The procedural road generation made it possible to present each participant with the same pool of parameterized turns in a randomized order for the purpose of reducing the influence of fatigue or experience on experimental results. Each trial consisted of a straight portion (20m), a curved portion of non-constant radius (100m), and a straight portion (20m). The trial ended when the participant exited the final 20m of straight road, at which time the preceding turn was removed and another curved portion of the road with its surrounding trees were added to the end of the roadway (two turns ahead), and the visual mask location was updated. This straight portion of the roadway also gave participants time to adjust their lane position and prepare for the upcoming turn over the span of a 40m straight-lane road segment. Because the participants speed was fixed at 19m/s, the total duration of a trial was roughly constant at 9.5 seconds.

Decisions regarding the turn radii that defined the curved portion of each road segment were based on rough estimates of the curvatures that would be acceptable in natural environments given estimates of the friction coefficients typical of a car’s tires on a dry day. On dry road conditions, a normal rubber tire demonstrates a coefficient of friction *µ* = 0.5 to 0.8 (Ghosh, 2011). The equation for minimum radius of curvature before skidding occurs is a function of speed, and is simply *R* = *v*^2^*/*(*µ* ∗ *g*), where *g* is gravity at 9.8 *m/s*^2^. Given our constant speed of 19m/s, this equation suggests that a reasonable range of radii that a car could follow without skidding would be 46m-74m. Although there was no skidding possible in this steering task, the radii of curvature were chosen within this range so as to maximize task difficulty but maintain reasonable radii of curvature. The decision to adopt non-constant radii of curvature was made to prevent the simple steering adjustments that could be used to navigate roads of constant curvature, and to instead encourage the online visually guided control of steering. The final chosen ranges were 46-56m and 66-76m, where the radius of curvature along the path increased from the first radius to the second at a linear rate over the course of the turn.

#### Gaze-contingent occlusion placement

The occluded quadrant was placed in one of four quadrants: upper left, upper right, lower left, or lower right. The mask was drawn using post-processing shader code, where certain pixels were replaced with a middle gray value after the scene had been rendered. The location of the gray pixels were determined according to the position of gaze on each frame. The Vive Pro Eye eye tracker provided gaze in world coordinates, and the data was converted to screen coordinates first to accommodate the post-processing effect. To block only one quadrant of the entire scene at a time, a rectangular area was defined before shifting the rectangle center up and left (in the case of the upper left occlusion) by one-half of the screen height and width. Foveal sparing with a radius of 2.5° was introduced because CB patients generally have a similar amount of spared central vision (Horton et al., 2021). This was implemented simply by only rendering pixels gray in the given quadrant if they exceeded the radius from the gaze location.

### Experiment design and procedure

The procedurally generated roadway was comprised of alternating straight and curved sections. Each trial (i.e., curve in the road between two straight-road segments) was parameterized by turn direction (left/right), turn radius (non-constant curvature at 46-56m or 66-76m), and the presence/location of the gaze contingent mask (upper left/right, lower left/right, or no occlusion), for a total of 20 possible trial types. Each trial was repeated eight times for a total of 160 trials, making the total experiment duration approximately 26 minutes. The order of trials was randomized for each participant.

Upon arriving at the laboratory, participants were informed of the purpose of the research and of the basic task design. The specific task instructions were to “steer naturally, while centering their head and body in the middle of the single-lane road”. This instruction was repeated twice, first during the initial explanation of the experiment, and then again just before they navigated their first turn. The task of staying as centered in the lane as possible provided participants with a goal to keep them engaged in an otherwise repetitive task, and it provided us with a reference for what would be considered “perfect” performance. A sample prerecorded video of the task was displayed for each subject to improve clarity of the task instructions. At this time, if participants wished to proceed, they signed the voluntary consent form and filled out a questionnaire with questions about demographics and driving and video game experience. Each person was then fitted with the headset. After completing the integrated Vive Pro Eye eye-tracking calibration sequence, the participant was transported to a stationary location at the start of the roadway and told that they could begin the task by clicking a button on the steering wheel. Participants were reminded that clicking this button mid-experiment would pause themselves if they needed to take a break due to VR sickness, in which case the headset could be removed and the eye tracker calibration repeated upon restart. Each subject practiced between 4-8 turns in the road to familiarize themselves with the wheel’s response in the virtual environment. The decision to begin the full experiment was made once participants appeared to navigate the turns comfortably. At this time, the task was reset, and the participant began the experiment.

### Steering analysis

For each participant, the effect of occlusion quadrant location on their lane position was calculated and compared to the steering behavior of the participant on control trials, in the absence of a mask. Lane position was measured at 90 Hz by taking the raw data of distance to the road center and subtracting the half width of the road (3 meters), yielding a measure of the participant’s distance from the inner road edge. Averages were computed across various segments of the turn, particularly the middle 40%, and the results could be separated by trial characteristics like turn direction, turn radii, and occluded quadrant condition.

### Gaze analysis

The estimated gaze direction vector within the head’s coordinate system was saved at 90Hz. Although the Vive Pro Eye eye tracker has the ability to return both the monocular and binocular gaze data, only the binocular signal was recorded and used to position the gaze-contingent masks in real-time. Prior to analysis, this gaze vector was subjected to a series of coordinate transformations. The raw gaze vector that described the direction of gaze within a head-fixed coordinate system was converted into the spherical coordinates of azimuth and elevation with respect to the vector protruding forward from the head in depth. The signal was then rotated to account for the additive effect of head orientation on gaze direction and transformed into a coordinate system in which azimuth and elevation were defined with respect to a static estimate of the participant’s natural “forward body position”. This position was sampled upon experiment start when the person was asked to put their hands on the steering wheel and gaze “straight ahead.” Hereafter, all discussion of gaze directions while driving will be considered within this default coordinate system. Because participants looked in the direction they were turning, the distribution of gaze azimuth samples was bimodal. To facilitate comparison of gaze behavior on left vs right turns of similar shape/turn radii, the gaze azimuth signal on left turns was negated prior to analysis.

To calculate the percentage of gaze samples that fell on the road while steering, gaze direction vectors below the horizon (i.e., with a negative elevation) were projected onto the ground plane. These points were then converted to world coordinates and filtered to identify those samples that fell within the boundaries of the road. Since the eye tracker introduced error into the estimate of the gaze direction (range of 0.47° −1.55°; discussed below), a threshold of *±*2° was added to assume that any gaze point within this threshold from the closest point on the road edge was a fixation on the road. These points were also used to calculate the linear “look-ahead distance” distance from the participant’s head to their gaze location on the upcoming road.

Saccade identification was performed using the gaze velocity signal, which was computed at each time step from the azimuth and elevation position and associated timestamps. Both velocity signals were filtered using rolling mean and median filters with kernel sizes of five, or 55 ms assuming a constant 90Hz. The following directionless velocity signal was computed from the filtered velocities: 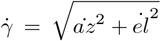. Saccades were identified from the directionless velocity signal where the velocity exceeded 30° per second. A reasonable minimum saccade time of 25 ms was implemented to prevent unwanted saccade identification from noise. Saccade landing points were also observed by identifying the fixations within a gaze signal and assigning the first gaze azimuth location of a fixation to be the saccade end point. Fixations were defined as the parts of the gaze signal where both the azimuth and elevation velocity signals were less than 20° per second. The first index where this condition was true was the beginning of a fixation and thus the saccade landing point. Saccade landing point distances were calculated by the distance from the gaze point to the head, and the previously-calculated gaze azimuth and elevation locations were also analyzed at each saccade landing point.

### Eye tracker calibration and accuracy assessment

Although the manufacturer of the Vive, HTC (VIVE, n.d.) claims to operate at 120 Hz with an accuracy of 0.5°-1.1° degrees, empirical measurements were taken with each participant following the short practice session with the steering task and prior to the start of the experiment. The custom calibration assessment involved the serial presentation of eight target dots 3 evenly arranged around a centered 10° invisible circle (see Figure 3). Only one target was visible at any time. Participants were asked to say “go” when they were looking at the target and ready to not blink for a second at a time. The researcher pressed a button to turn it yellow, during which the gaze position was recorded to a .csv tracker file for the duration of a second. The recording at a particular location was repeated if the participant blinked. Gaze position was recorded first at the central target, located at (0,0) in azimuth and elevation relative to the head, then next at all eight surrounding targets in a counterclockwise direction beginning with the point right of center. Pilot testing suggested that the azimuthal component of gaze covers a wider range of the visual field than the vertical direction when performing the steering task. Consequently, the assessment grid was extended to include assessment targets at 5°, 10°, 15°, and 20° to the right and left of screen center.

The spherical coordinates of the binocular eye-in-head vector were calculated by taking the arctangent of the raw gaze direction data x/z (azimuth) and y/z (elevation), and converting the result to degrees. The ground truth values of gaze during each one-second interval recording were logged relative to (0,0) which represented the “straight-forward” orientation of the head, centered in the headset’s field of view. At each calibration point, the standard deviations (*σ*) of all gaze samples in azimuth and elevation were computed, and the horizontal and vertical 95% confidence intervals were defined as 2 ∗ 1.96 ∗ *σ*_*az*_ and 2 ∗ 1.96 ∗ *σ*_*el*_. Gaze error was calculated per target location in the azimuth and elevation dimensions (mean of the absolute value of gaze samples at target minus target value), and these errors were combined into the effective gaze error via 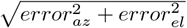 (units of degrees). To find the final gaze error per participant, gaze errors at all target locations were averaged. Gaze errors ranged from 0.47° at best to 1.55° at worst (see Figure 3 for examples).

### Eye tracker latency

The Vive Pro Eye was selected for this study based on its compatibility with our experimental setup and its established performance metrics. Although more recent measurements of the device’s end-to-end latency (79–81 ms) (Stein et al., 2021) place the Vive Pro Eye at the upper end of the optimal range suggested for gaze-contingent effects (50–70 ms) (Albert, Patney, Luebke, & Kim, 2017), both our CPU and GPU were notably more powerful than the hardware featured in (Stein et al., 2021) (CPU comparison: AMD Ryzen 9 5950X 16-core processor (ours) vs. Intel Core i9 9900K (3.6 GHz), GPU comparison: NVIDIA GeForce RTX 3080 Ti (ours) vs. NVIDIA RTX 2080). CPUs and GPUs with greater power are more likely to increase VR performance capabilities including reducing latency (Tytarenko, 2023). Participant feedback indicated that they were unbothered by the gaze-contingent manipulation, and that it had an adequate level of temporal responsiveness needed to complete the task naturally. Participants were also asked to report whether they could momentarily fixate within the gray occlusion area – a task theoretically impossible under conditions of low latency. No participant reported being able to fixate within the occlusion after the eye tracker calibration or during the experiment when asked, suggesting that the latency was not perceptibly intrusive for our task. We concluded that although there are almost certainly subtle and unintended effects related to the use of a gaze-contingent display with non-negligible amounts of latency that limit the Vive Pro Eye’s utility in more stringent gaze-contingent paradigms, its performance in our context was sufficient to achieve the study’s goals while maintaining participant comfort and experimental feasibility.

### Statistical analysis

Two repeated measures ANOVAs were conducted on the steering data with the independent variables of turn direction, turn radius, and occlusion quadrant. Lane position and wheel variance data were the dependent variables, and post hoc comparisons with Holm-adjusted p values were used to compare means of groups within the significant effects. The assumption of sphericity was tested for all ANOVAs in this study and sphericity corrections were applied where necessary using the Greenhouse-Geisser method. All results were reported with Greenhouse-Geisser correction applied (where sphericity was violated).

To directly test the steering symmetry, or lack thereof, between left and right visual field occlusions, the lane position and wheel variance data were relabeled by participant according to the side on which the occlusion fell, and paired t-tests were evaluated. These t-tests were executed to identify whether having a left or right visual occlusion significantly affected driver lane position or wheel variance. Assumptions of normality and equality of variance were checked for each t-test and corrected if necessary using log transformations and Welch’s t-test to address variance equality violations.

Gaze data was analyzed with multiple ANOVAs, all with turn direction, turn radius, and occlusion quadrant as independent variables. The following gaze metrics were evaluated with an ANOVA: saccade frequency, gaze azimuth, gaze elevation, average percent of gaze on road, and average look-ahead distance. Additional statistics were computed on the subset of saccade data in which participants made saccades towards an occlusion when it was present. Four paired t-tests were used to compare control saccade data versus mask data in each of the four quadrants, with a Bonferroni-corrected alpha of 0.0125 to account for multiple comparisons. One additional paired t-test was conducted after regrouping the data based on whether the occlusion was on the left vs. right of the visual field. This test made it possible to investigate the potential of asymmetric differences between left vs. right visual field occlusions. The t-test statistical analyses on saccade frequency, magnitude, and direction were repeated in the follow-up study with eight additional participants who were subjected to only an upper left or upper right quadrant during the experiment.

## Results

The trajectories of participants over the course of a trial revealed distinct differences in steering specifically in the presence of gaze-contingent occlusions in the upper visual field. Figure 4 depicts the average trajectories along the curves from trial start (t=0) to trial end (t≈7.3) for all 24 participants, separated by turn radius and occlusion quadrant. As participants increased their corner cutting, their position shifted to the left of the vertical dotted line in Figure 4 that marks the road center. Thus, the data showed that all participants corner cut, for almost the entire span of the trial. Because three of the trajectories (no mask, lower left, and lower right occlusions) are positioned further to the left on the x axis for both turn radii conditions, the data highlight the fact that upper visual field occlusions decreased corner cutting. This effect was exaggerated in the central part of the trials, on the curved portion of the road specifically, and for this reason all further analysis is focused on the middle 40% (from ≈2.2-5.1 seconds).

A central goal of this study was to test for steering biases that were dependent on mask location, and Figure 5 confirmed that imposing an artificial blind field in a quarter of the visual field affected steering differentially, depending on the mask location. Figure 5 plots the average lane position during the middle 40% of trials for each occlusion condition relative to the control condition in which no mask was present (black dotted line). In order to compare the effect of the mask on steering behavior within subjects, the within-subject average lane position on left and right turns was subtracted from all other left and right turns within-subject before analysis. Statistical analysis revealed a main effect of occlusion quadrant on steering 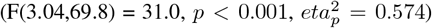, and an interaction of turn direction with occlusion quadrant on steering performance 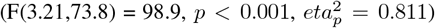. The interaction of turn direction and mask location is most easily interpreted when directions are considered relative to the mask location (e.g. turns towards vs away from the mask), as presented in Figure 5. The figure illustrates a consistent reduction in corner cutting when participants turn towards a mask (blue dots) compared to their steering in the control condition (dotted black line). This observation was confirmed with pairwise comparisons between the trials in which participants turned *towards* a mask and the trials where they turned in the same direction without a mask (e.g., turning left with a mask in the upper left vs. turning left in the absence of a mask); all four such comparisons reached statistical significance at the *p <* 0.001 level. When turning *away* from an occlusion in the lower half of the visual field, corner cutting was increased relative to the control condition (lower left turning right: *p <* 0.001, lower right turning left *p <* 0.001), but there was no difference when the mask was in the upper half of the visual field.

Qualitatively, gaze behavior involved the tracking of points that were stable in the virtual environment (but moving through the visual field of the translating observer), followed by saccades in the opposite direction in the direction of the turn they were actively navigating (i.e., optokinetic response and nystagmus). This behavior is shown in Figure 2B, and similar to the behavior that has previously been characterized in two studies that featured a similar task environment (Giguere et al., 2024, 2025). This characterization is also present in Figure 6, which displays the frequencies and directions of saccades made during the middle 40% of the turns. The figure shows that saccades were generally made towards the upper right quadrant when navigating right turns and towards the upper left quadrant when navigating left turns, regardless of the occlusion location. The presence of a mask disrupted this behavior and reduced overall saccade frequency 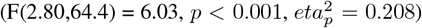. This is most notable in Figure 6 through the interaction of mask location and turn direction 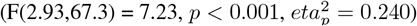. Post hoc pairwise comparisons revealed that saccade frequency for the control condition differed significantly from all masked conditions except for the lower left (lower left vs. control: *p* = 0.072, lower right vs. control: *p* = 0.034, upper left vs. control: *p* = 0.005, upper right vs. control: *p <* 0.001) indicating that the presence of a mask often reduced saccade frequency by about 3.5 saccades across the middle 40% of the turns. Similarly, pairwise comparisons of the interaction between quadrant and turn direction on frequency revealed that turns towards an upper visual field occlusion reduced saccade frequency relative to the control (upper left vs. control turning left: *p* = 0.019, upper right vs. control turning right: *p <* 0.001, while for the lower visual field occlusions, only turns away from a lower left mask differed significantly from the control (lower left vs. control turning right: *p* = 0.017). In summary, the data suggest that the presence of a mask reduced saccade frequency, especially when an observer was turning in the direction of a mask obscuring the upper visual field.

**Figure 2:**
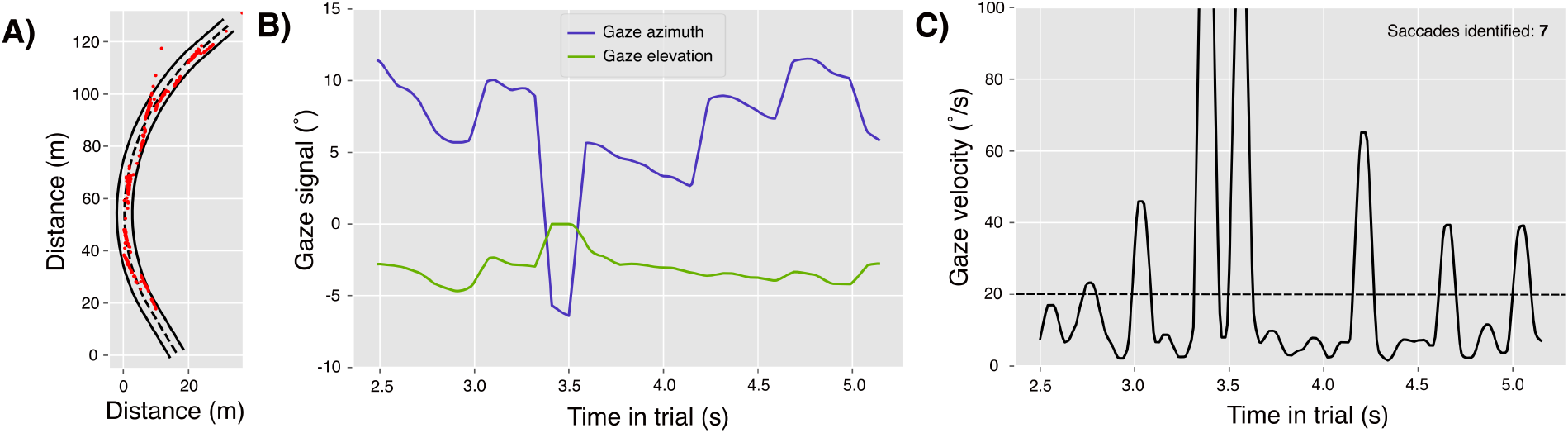
Gaze data from a single trial. **2A)** Top down view of the road from a single trial. The turn begins at the bottom of the panel and ends at the top, where the first 20 meters is straight road, followed by a curve with a non-constant turn radius that changes from 66m to 76m, then the last 20 meters of straight road. The red dots indicate samples from the participant’s gaze direction when their gaze intersected with the ground plane. Because participant’s looked ahead of their current position, there are no red dots present on the first section of the road. **2B)** Gaze azimuth and elevation signals. Elevation is negative because zero degrees elevation is the horizon. Azimuth varies relative to zero degrees which is defined as the “forward body direction” that was sampled upon experiment start. **2C)** Gaze velocity signal derived from the combined azimuth and elevation signals 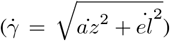. The horizontal dotted black line indicates the saccade threshold used for saccade identification. The detector identified six saccades on this trial.

**Figure 3:**
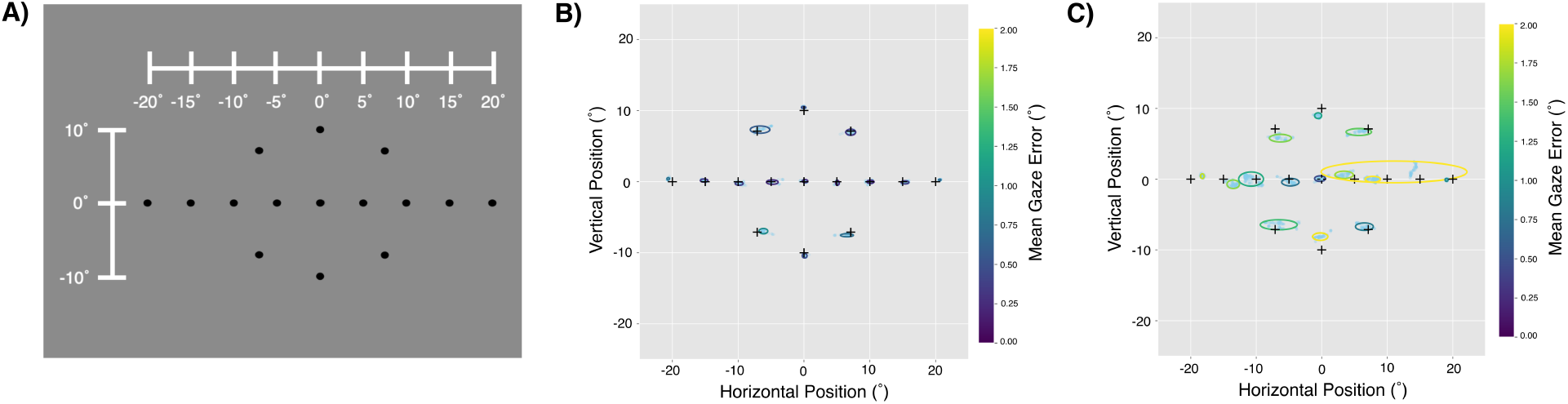
**3A)** Calibration assessment points as viewed from inside the headset. Field of view is cropped for visualization purposes. The white scale above the black dots was not visible inside the headset and depicts the azimuth and elevation scales according to the participant. Only one black dot at a time was visible during calibration assessment. **3B)** Calibration assessment grid for the participant with the lowest average eye tracking error (0.47°). Each cross marks the location of the calibration assessment point in degrees azimuth and elevation. The light blue dots represent all gaze samples collected during each 1 second of recording at each of the cross locations. The ellipses are color coded according to the color bar where the edge color corresponded to the mean gaze error at each point. The ellipse shape is defined by the 95% sample confidence intervals in both the vertical and horizontal directions. **3C)** Calibration assessment grid for the participant with the highest average eye tracking error (1.55°). All metrics are the same as described in Figure **3B**).

**Figure 4:**
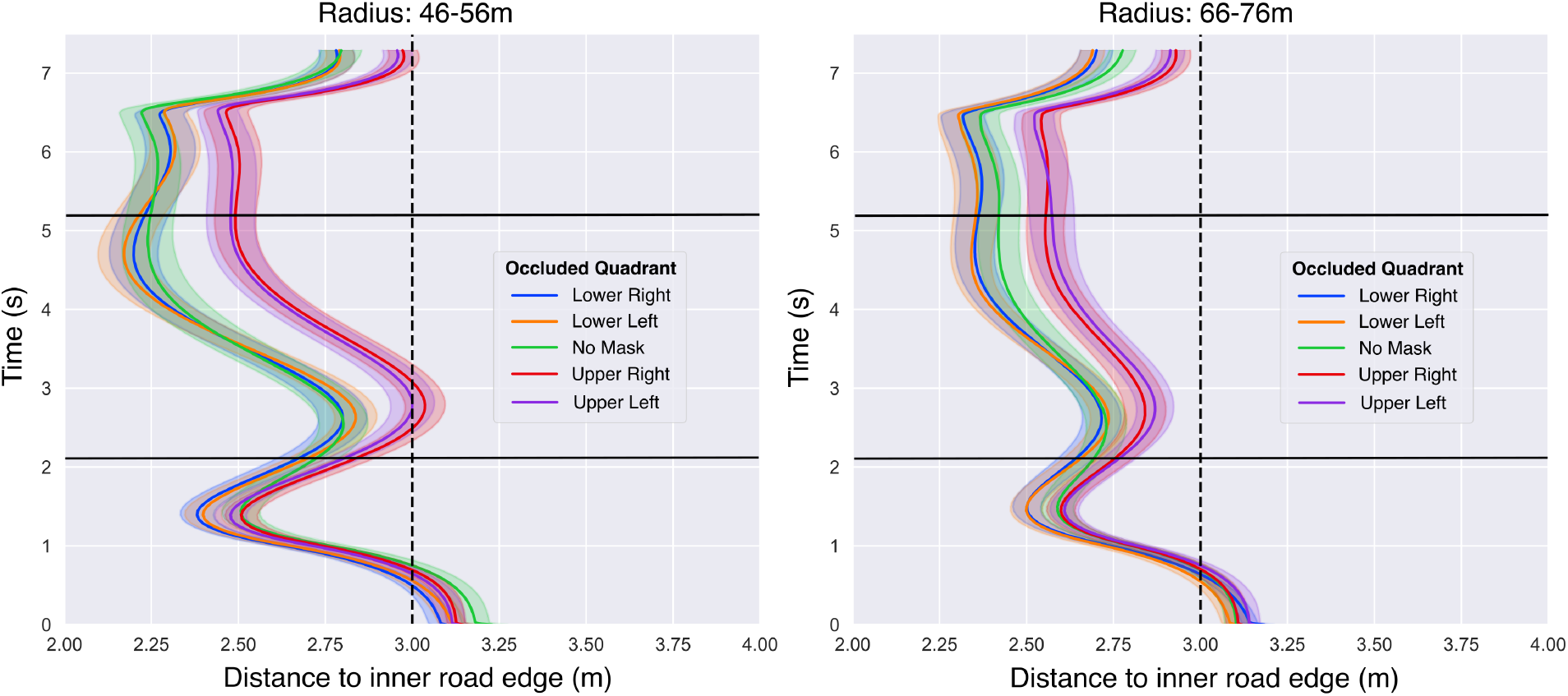
Average lane position over time for the two turn types of non-constant radii, separated by occlusion quadrant. Trials with a sharper turn radius are plotted on the left and the wider turn radii are on the right. Although the entire lane width was 6m, these figures display only the area spanning the central third of the road width. Averages of behavior while navigating turns are computed over the central 40% of the turn length (indicated by the black horizontal lines), which is also the portion over which differences in lane position were the most exaggerated and stable. The vertical dotted black line marks the center of the road. Lane positions left of the vertical line indicate a lane position that is biased towards the inner road edge, which we refer to as “corner cutting”. The trajectories navigated in the presence of an upper-field occlusion (purple and red lines) demonstrate less corner cutting compared to trials with a lower-field occlusion (blue and orange lines) and the control (green line).

Because these results indicated unique gaze behavior when making saccades in the direction of an upper visual field occlusion, Figure 7 plots the average magnitude and directions of saccades made in the direction of the masks when present (blue dots). This data is compared to the control condition, where no mask was present but saccades were made to each of the four quadrants (red dots). From Figure 7, it is evident that the presence of an occlusion mask impacts the magnitude, but not direction, of saccades made towards the upper visual field. This observation is confirmed by four paired t-tests between control and mask conditions for each of the four quadrants (with a Bonferroni correction of *α* = 0.0125), which revealed significant differences between the control and masks in the the upper visual field, but not the lower (upper left vs. control: *p <* 0.001, upper right vs. control: *p <* 0.001, lower left vs. control: *p* = 0.070, lower right vs. control: *p* = 0.041).

In our main experiment, the between-trial/turn randomization of mask location quadrant gave participants little time to learn gaze strategies to compensate. To explore the potential effect of learning over slightly longer periods of time, the study was replicated with a blocked-design instead, where eight subjects were tasked with steering in the presence of either an upper left or an upper right visual occlusion. Participants navigated 16 control trials first with no occlusion mask, 32 trials with the mask, then 16 trials without the mask once again. Figure 8 displays the results. The distribution of saccades as well as saccade magnitudes and directions are quite similar to the data represented in Figures 6 and 7. Despite the visual similarities, paired t-tests on saccade frequency between control and occlusion conditions, saccade magnitude, and saccade direction did not reveal significance where the data from the main study did. We suspect that this can be attributed to low statistical power from only eight participants rather than the 24 of the main study. Additionally, we separated the follow-up data by trial type (according to mask (on/off), turn direction (left/right), and turn radius (46-56/66-76)), and plotted saccade magnitudes and directions over time. Data from each participant was observed independently, and there were no clear trends to indicate a learning effect over time. With the caveat that this follow-up study had limited participants, we did not find any evidence that gaze strategies change over time in response to a fixed visual occlusion.

The smoothness of steering movements was measured as the variance of wheel orientation within the central 40% of the turn (Figure 9). A repeated measures ANOVA revealed significant effects of quadrant location 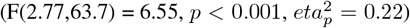, turn direction 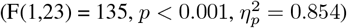, and their interaction 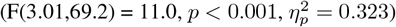. Visual inspection of Figure 9 indicates that variance differed from the control condition only when making turns away from the mask when it was located in the lower visual field (paired comparisons for right turns with a lower-left mask vs. right turn no mask: *p <* 0.001, left turn lower-right mask vs. left turn no mask: *p* = 0.004).

Although we chose to vary turn radius (46-56m or 66-76m) solely for the purpose of increasing task difficult and requiring real-time steering visuomotor adjustments, it is worth mentioning that radius also affected steering and gaze behavior. The interaction of occlusion quadrant and turn radius impacted average distance from the inner road edge 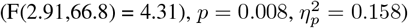, as did turn radius on the saccade frequency 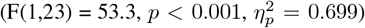 and wheel variance 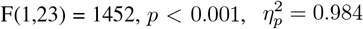. Their significance indicates that the sharpness of a turn influences steering and gaze behavior.

## Discussion

This virtual steering task was designed to test the hypothesis that the impact of cortical blindness on natural steering behavior is most consistent with an omission/occlusion of visual information, rather than a disruption to the cortical processing of task-relevant visual information. Visually-healthy participants were tasked with driving a virtual reality roadway while a gaze-contingent occluding mask was imposed on a quadrant of their visual field. We reasoned that if a simple visual occlusion is sufficient to explain the biases observed in the presence of CB, then the visually-healthy drivers in this study would demonstrate an identical pattern of biases as has been observed previously when CB drivers participated in the same task. However, the data produced by this study indicate that visually-healthy participants subjected to gaze-contingent masks did *not* demonstrate the same pattern of steering biases previously observed in CB drivers. On this basis, we conclude that the pattern of steering biases observed in CB likely arise because of a disruption to cortical processing, learned compensatory behaviors, or a combination of both. More broadly, we conclude that the effects of cortical blindness on steering behavior may not be consistent with a simple occlusion or omission of visual information.

Although the visually-healthy participants in our study did not demonstrate the same pattern of biases as those with CB, they did demonstrate biases that provide new insight into how visual information is used to guide steering. The presence of a mask in any location decreased corner-cutting on turns towards the mask, but only occlusions of the lower visual field caused an increase in corner-cutting when turning away (Figure 5). A lower visual field occlusion tends to block part of the near road. Because the visible near road facilitates an increase in lane accuracy at the cost of more variable adjustments of the steering wheel (Salvucci & Gray, 2004; Land & Horwood, 1995; Donges, 1978; Lappi, 2014; Frissen & Mars, 2014; Mole, Kountouriotis, Billington, & Wilkie, 2016), one might expect an occlusion of the lower visual field to decrease the participant’s wheel variance because they might rely instead on far road information. However, Figure 9 shows that wheel variance *increases* on turns away from lower visual field occlusions, and does not change on turns towards them. The results appear inconsistent with the theories proposed by previous researchers (Salvucci & Gray, 2004; Mole et al., 2016; Land & Horwood, 1995), but there may be an alternative explanation. Only part of the near road is occluded with the simulated masks in this study, so it is not entirely correct to suggest that the near road information is missing. In the case of the lower visual field occlusions, participants could be relying on the far road in addition to the part of the near road that is still visible. A more likely reason for the increase in wheel variance on turns away from lower visual occlusions may be related to the relative visibility of the road edges. According to Figure 1, the near-road outer road edge is obscured from view only when turning away from a lower visual occlusion. In contrast, when the near-road outer road edge is visible for turns away from an occlusion in the upper visual field, steering remains *unchanged* (Fig. 5) relative to control conditions. Thus, if participants are in part relying on road edge information to navigate, it could explain why corner cutting and wheel variance increased when turning away from a lower, but not upper, visual field deficit.

**Figure 5:**
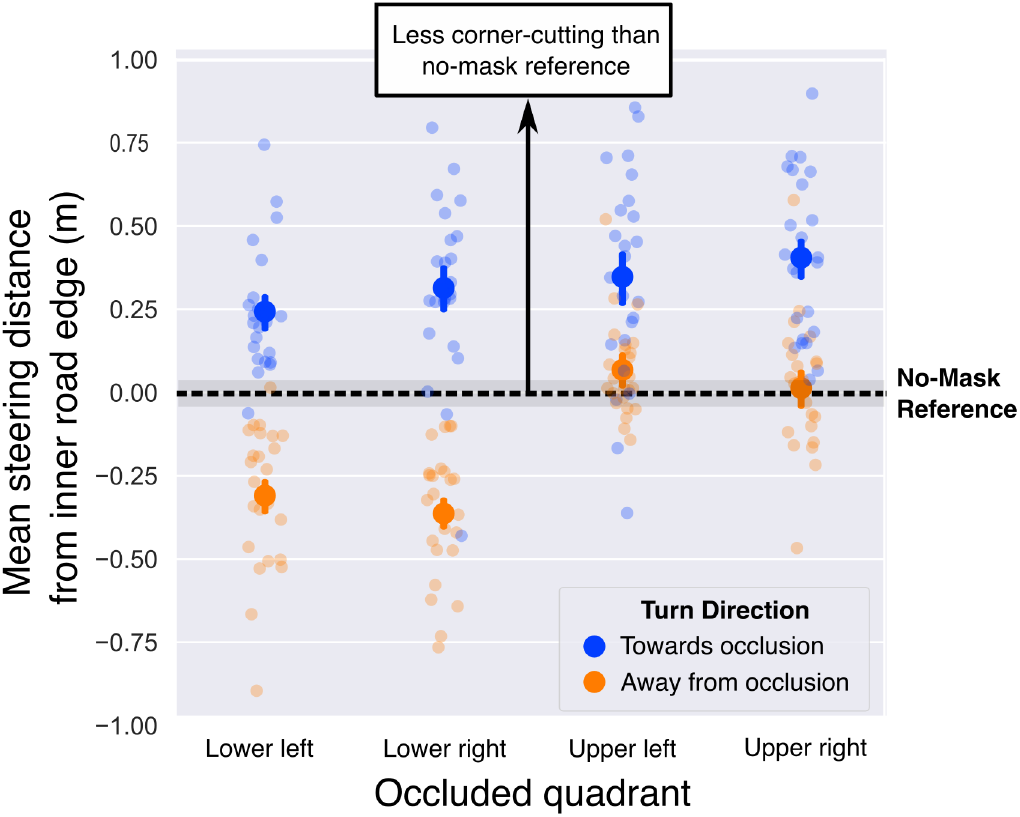
Mean steering distance from the inner road edge separated by turn direction and occlusion quadrant conditions for all participants. All data points shown are in reference to the mean distance from the inner road edge that each participant exhibited on the no-mask control trials for left and right turns. The data is coded by whether the turn was towards or away from the occlusion. The black horizontal line indicates the reference, such that if a point were to lie on the dotted line it would suggest no steering difference relative to performance in the control condition. The shaded gray region around the horizontal black line indicates the average within-subject error on the control trials. Points above the horizontal dotted line represent less corner-cutting than the control condition, and points below indicate more. Error bars indicate the 95% confidence interval of within-subject error.

**Figure 6:**
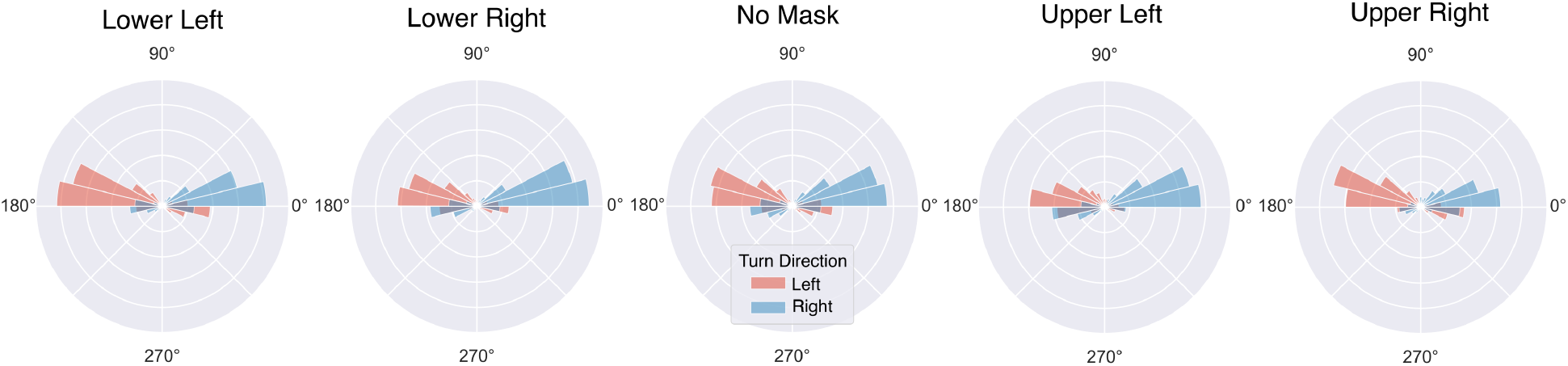
Normalized polar histograms of pooled saccades from all participants over the middle 40% of each trial, separated by occlusion quadrant condition. All subplots are normalized to the same arbitrary maximum count that scales all data to within 0-1, such that the frequencies across subplots can be directly compared to each other. The direction of the saccade is represented by theta, where 0° would correspond to a saccade directly to the right of fixation, and 90° would be a saccade directly upward. Red bins indicate saccades made when navigating left turns while blue bins denote the saccades made on right turns. The figure illustrates how most saccades during the middle 40% were made upward, and generally in the direction of the turn.

There is some related evidence from previous research to support the interpretation that the visibility of road edges can influence steering. One study looking at driver gaze in a virtual environment while varying the visibility of inner and outer road edges (fully opaque, faded, or removed) found that people give more weight to more reliable sources of visual information (Kountouriotis, Floyd, Gardner, Merat, & Wilkie, 2012). This finding is supported by research on visual cue combination (Ernst & Banks, 2002; Landy, Maloney, Johnston, & Young, 1995; Newman, Qi, Mou, & McNamara, 2023; Chen, McNamara, Kelly, & Wolbers, 2017). Kountouriotis et al. removed either the inner or outer road edge in a virtual steering task and observed participant’s lane positioning relative to when both road edges were present. Although they reported no significant differences in lane position when removing one road edge at a time, their task instructions required participants to maintain a defined fixed road position that was indicated at the beginning of each trial, whereas our participants were instructed to maintain a central lane position only at the beginning of the study. As a result, participants could be biased in lane position towards the visible road edge which would be a natural response to the uncertainty introduced when the opposite road edge is occluded and when the target lane position is not visually marked upon trial start. Alternatively, lane bias towards towards the visible road edge could also reflect a shift in strategy from trying to stay roughly centered in the lane to a strategy of maintaining a fixed distance from the visible road edge. For these reasons, if one road edge is blocked by a simulated occlusion and people rely more on the visible road edge, it is reasonable to expect increased corner-cutting when only the inner road is visible and decreased corner-cutting when only the outer road edge is visible. This is consistent with our results: when occlusions blocked the inner road edge, people corner-cut less, and when they blocked the outer road edge, people corner cut more (see Fig. 5). The *only* cases in which corner-cutting remained unchanged relative to the control were also the only cases in which both road edges were almost entirely visible (turning away from upper left/right quadrants).

The observation that people were biased in lane position towards the visible road edge can be compared to the steering of CB participants with gaze-contingent visual deficits, some of whom are biased in lane position away from their blind fields and towards the more visible road edge. If visually-intact participants corner-cut less on the side of a simulated visual field deficit when turning towards it (as shown in Fig. 5), then it is not surprising that some CB patients exhibit that behavior as well. However, because previous data has revealed that those with left-sided CB are biased towards their blind fields but those with right-sided CB are not (Giguere et al., 2025), and visually-healthy participants are biased towards the visible road edge equally regardless of an occlusion on the left or right, these results imply that perhaps cortical damage affecting the left visual field affects steering more like a visual occlusion than cortical damage affecting the right visual field. It is also possible that a combination of factors like CB disruption of cortical processing and their learned compensatory behaviors contribute to the different steering biases seen in left and right-sided CB participants.

This study was also designed to address whether the presence of a gaze-contingent occlusion mask would change the gaze patterns of participants while steering and whether the location of the occlusion was particularly influential. We found that gaze was similar between quadrants conditions with a few exceptions. In all cases, participants tended to saccade towards the upper left quadrant when turning left, and the upper right quadrant when turning right. This pattern could be explained as a reflection of the gaze behavior necessary to complete the steering task successfully. Participants also made fewer saccades in the direction of the occlusion when it was placed in the upper visual field, and these saccade magnitudes were smaller relative to the saccade magnitudes in the same direction without an occlusion present (Fig. 7). Because the saccade magnitudes towards an occlusion exceeded the dotted circle in Figure 7 that marks the edge of the visible field of view, they were large enough to land inside the area of the mask where it was prior to the saccade. Given that saccade direction remained unchanged, this behavior can be interpreted as a moderate change in gaze compared to the control condition, and it might reflect participant uncertainty regarding the lack of visibility of the visual environment behind the mask. A saccade of full magnitude, such as the ones measured when there is no visual occlusion, could be disorienting if attempted in the presence of an occlusion. In other words, our results provide evidence that a gaze-contingent occlusion in the upper visual field encourages more conservative gaze behavior in the form of smaller saccades and lower saccade frequency.

**Figure 7:**
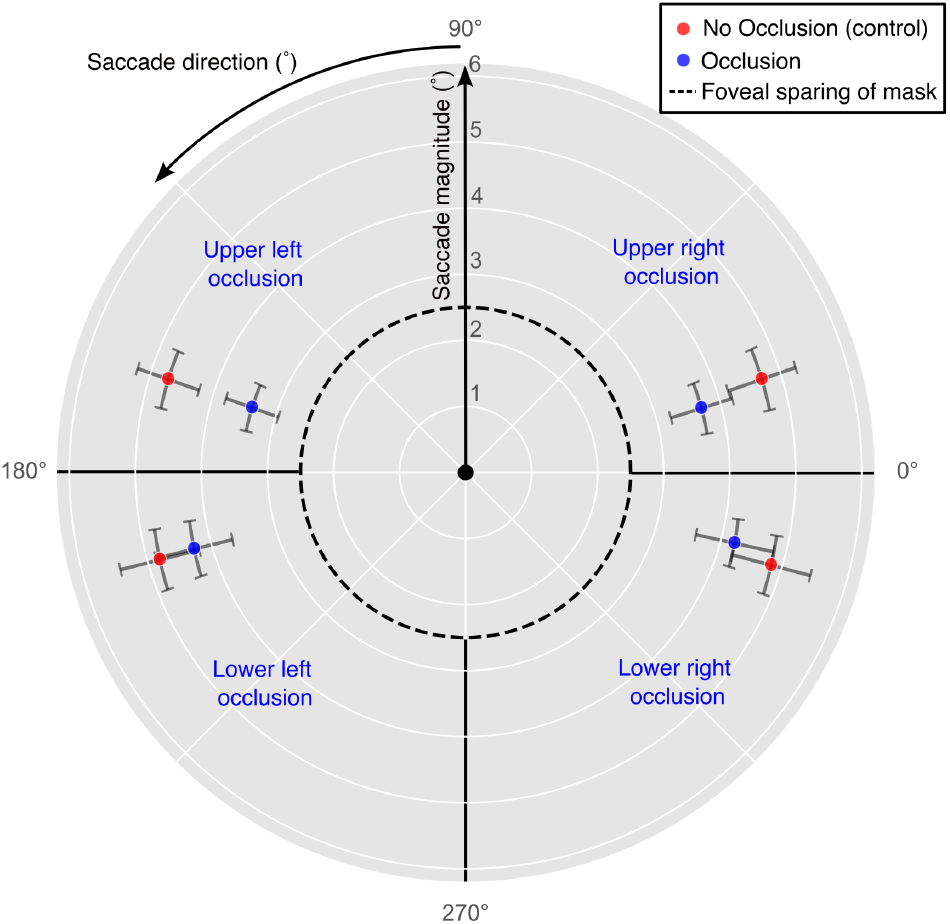
Polar plot of the saccades made towards an occlusion during the middle 40% of each trial. For example, the data represented in the upper right quadrant include only the saccades made to the upper right in the presence of an upper right visual occlusion (blue dot), or the saccades made to the upper right in the control condition (red dot). The average magnitude of saccades under each condition is represented by each dot’s distance from the center of the figure, where all saccades begin. The average direction of saccades is denoted by the angle theta, where 0° is a saccade directly to the right of fixation, and 90° is a saccade directly upward. Error bars are marked as the 95% confidence intervals in both the magnitude and direction of saccades. The confidence intervals reveal a significant effect of saccade magnitude in the upper visual field only, while saccade direction is insignificant in all four quadrants.

**Figure 8:**
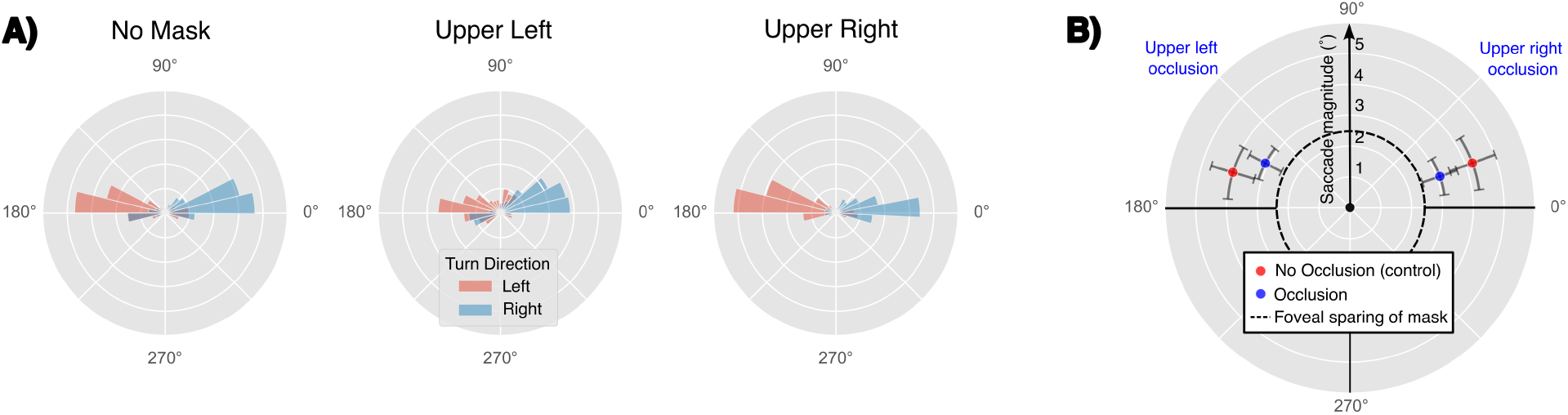
Summary of the data from the follow-up study with eight additional participants steering first without an occlusion (16 trials), then with an occlusion in a fixed location (32 trials), then without the occlusion once more (16 trials). **8A)** Polar frequency histograms of all saccades within the middle 40% of trials. The data from four participants each are depicted in the upper left and upper right frequency histograms respectively, while all control data from the eight subjects is pooled in the control (no mask) case. Because there were double the amount of control trials than occlusion trials, the frequencies are scaled by 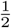 in the control histogram to promote direct comparison to the frequencies in the masked-data histograms. Each histogram is normalized from 0 to 1 on the same scale, where the outermost circle represents maximum counts. Saccade direction is indicated by the angle, where 0° represents a saccade direction to the right of fixation, and the bins are color coded by whether the saccades were made on left turns (red) or right turns (blue). Visually, saccade frequency towards the quadrant of the visual occlusion appear reduced relative to the control. **8B)** Polar plot indicating the average saccade magnitudes and directions of saccades made to the upper visual field when no mask was present (red dots), and when the mask was present (blue dots). The blue dots include only the saccades made towards the quadrant of the visual occlusion and include data from four participants each, while the control data combines saccades from all eight participants that were towards towards either the upper left or upper right.

**Figure 9:**
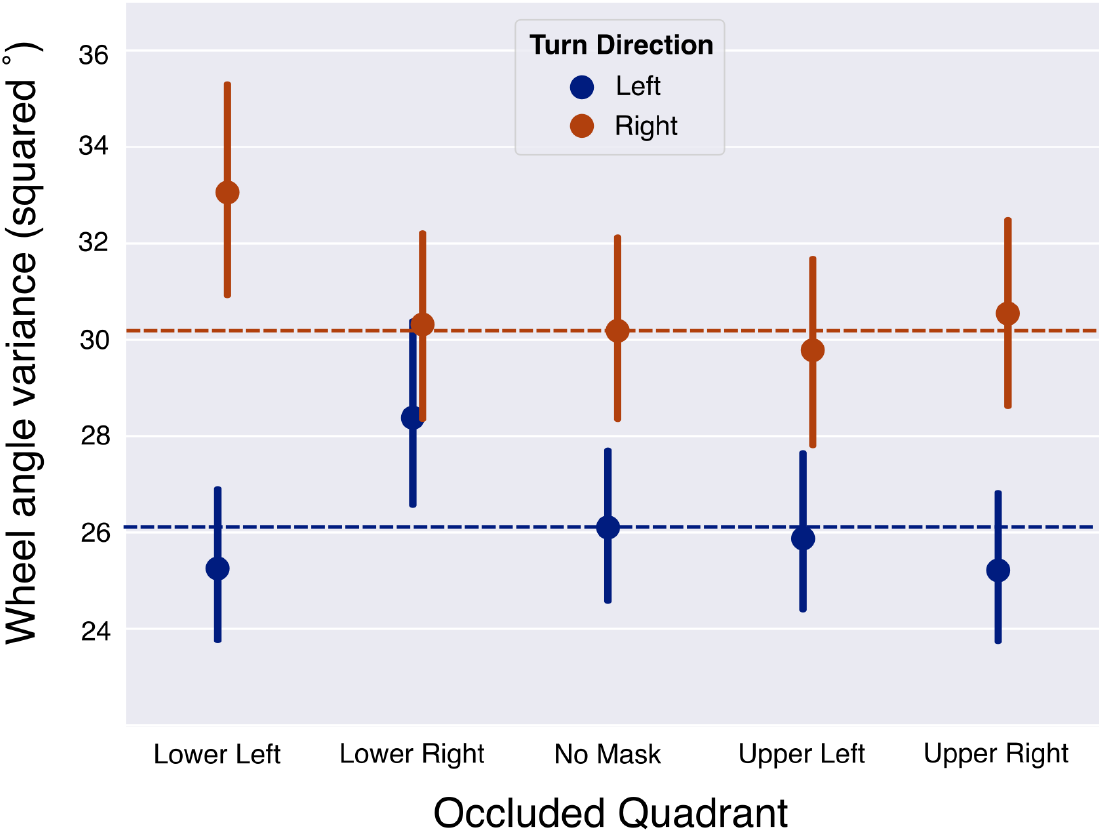
Steering wheel variance separated by turn direction and occlusion quadrant. The horizontal dotted lines represent the means of the reference conditions to which all other points of the same color can be compared. The plot reveals that wheel variance increases relative to the control conditions only when turning away from the blind field with a lower visual field simulated deficit. Error bars indicate the 95% confidence interval of within-subject error.

Furthermore, when saccade magnitude decreases, peak velocity also decreases, raising the potential concern that saccades may exist below our chosen threshold of 20°*/sec* which would effectively reduce frequency and yield a misguided result. The fact that we found a decrease in saccade frequency for upper visual field occlusions as well as a decrease in saccade magnitude prompted an investigation into this potentially misleading result. However, rerunning the analysis with a threshold of 15°*/sec* did not change the conclusion, and lowering the threshold to 10°*/sec* infringed upon the velocity for slow-moving fixations. This reduced the likelihood that saccades of lower peak velocity were undetected by our algorithm, and strengthened our interpretation that the decrease in saccade frequency towards upper visual field occlusions is a response to the occlusion.

Analysis of the non-saccadic gaze signal segments did not reveal any additional insights into how gaze behavior may change according to occlusion quadrant location, but the comparison of gaze and steering behavior together may provide evidence of compensatory strategies adopted when visual occlusions are present. Overall, the gaze results can be summarized by the findings that saccade frequency and magnitude are reduced when directed towards occlusions in the upper visual field locations. Because this steering task required saccadic eye movements towards the upper visual field, these occlusion locations were naturally more disruptive to the otherwise normal gaze behavior. However, while saccade magnitude and frequency were positively correlated with steering bias relative to the control (see Fig. 5, reduced saccade magnitude/frequency corresponded to reduced corner-cutting in the presence of upper visual field occlusions), participants also cut corners less when turning in the direction of lower visual field occlusions despite there being no change in saccade magnitude or frequency under these conditions. Because the upper visual field occlusions are located directly in the line of natural saccadic behavior while navigating the curves, the reduction in saccade magnitude and frequency could be interpreted as a compensatory mechanism to preserve steering behavior as much as possible. Because only a small percentage of saccades are made towards the lower visual field areas, it would not be necessary for gaze behavior to change in order to achieve a desired lane position. These results imply that occlusion of visual information in the upper visual field is more likely to alter gaze patterns with the effect of achieving similar steering performance to the steering under lower visual field occlusion conditions.

Lastly, our follow-up study investigated potential adaption to the visual occlusion, and it did not reveal any learning effects or apparent changes in saccade distribution. While this follow-up study was useful in promoting an assessment of whether gaze strategies changed when the occlusion location was fixed, it had limitations regarding its generalization to the “blind fields” in individuals with CB. Patients with CB often have lived with their visual deficit for months prior to participating in research studies, while the participants in the present study only had about ten minutes to adapt to the occlusion. Humans are naturally very good at adapting to changing environmental conditions as is evident by our ability to respond to changes in luminance from weather when driving for example, or our ability to quickly recalibrate to the sensitivity of a brake pedal in a new car, but adapting to CB may be a more involved process that takes more time. The present study involved no risks such as pedestrians, other cars, or traffic lights, thus successful completion of the experiment did not require frequent saccades to locations far in the periphery or even towards the occlusion in some cases. Ultimately, our findings that participant gaze behavior was similar when the occlusion location was fixed versus randomly placed motivates the need for future work with a more involved driving simulation requiring a larger variety of eye movements to see if the similarities persist. Such a task could potentially highlight learned gaze behavior even in visually-intact participants if faced with multiple close collision experiences or nearly-missed red lights.

## Conclusions

The results of this study indicate that the effect of CB on steering behavior should be characterized as either a disruption to cortical processing, a learned compensatory behavior, or a combination of both. By comparing the steering behaviors of visually-healthy participants with gaze-contingent occlusions to the steering of individuals with CB, we found that the latter exhibit steering biases that cannot be fully explained by visual occlusion alone. Our findings highlight the potential role of road edge visibility in steering control, reinforcing the idea that visually-healthy drivers dynamically adjust their behavior based on the most reliable visual cues available. Importantly, gaze analysis revealed that occlusions in the upper visual field prompted reductions in saccade frequency and magnitude, suggesting a potential adaptive mechanism for maintaining reasonable steering performance. These results contribute to a broader understanding of how visual information is integrated during steering. Further studies could explore how adaptive gaze and steering behaviors develop over time in CB patients and whether targeted interventions can facilitate appropriate compensation for such visual impairments when steering.

## Acknowledgments

This research was supported by the Research to Prevent Blindness/Lions Clubs International Foundation Low Vision Research Award (LVRA). We would like to thank Sijal Jaradat for his help in the programming of the Unity driving simulation environment.

## Footnotes

1 In Giguere et al. (2025), the effect of turn direction and group (e.g. controls, left CB, right CB) on lane position was significant. Post-hoc paired comparisons of within-group effects demonstrated significance for controls and right CBs only, however, all between-group comparisons within the same turn direction were insignificant. This likely reflected the fact that post-hoc paired comparisons are conservative and are less likely to reach significance especially with Bonferroni-corrected p-values and smaller sample sizes, which are typical in studies involving between-group comparisons of special populations (Kim, 2015). Visual inspection of the data separated by group and turn direction supports the interpretation that left CB are biased in lane position away from their blind fields.

2 Although visually-healthy drivers are also biased in lane position away from other close-by or moving vehicles (Delpiano, 2021), we note that the present study investigated steering only in the absence of other vehicles and moving obstacles.

## Notes

### Competing Interest Statement

The authors have declared no competing interest.

## References

Albert, R., Patney, A., Luebke, D., & Kim, J. (2017). Latency requirements for foveated rendering in virtual reality. ACM Transactions on Applied Perception, 14(4), 1–13. https://dl.acm.org/doi/10.1145/3127589, doi:10.1145/3127589 [Article]

Alipour, A., & Kazemi, S. (2015). Dominant right hemisphere controls biased rotation perception. Psychology & Neuroscience, 8(4), 435–441. https://doi.apa.org/doi/10.1037/pne0000034, doi:10.1037/pne0000034 [Article]

Bahnemann, M., Hamel, J., De Beukelaer, S., Ohl, S., Kehrer, S., Audebert, H., et al. (2015). Compensatory eye and head movements of patients with homonymous hemianopia in the naturalistic setting of a driving simulation. Journal of Neurology, 262(2), 316–325. http://link.springer.com/10.1007/s00415-014-7554-x, doi:10.1007/s00415-014-7554-x [Article]

Barthélémy, S., & Boulinguez, P. (2002). Manual asymmetries in the directional coding of reaching: further evidence for hemispatial effects and right hemisphere dominance for movement planning. Experimental Brain Research, 147(3), 305–312. Available from 2025-01-08 http://link.springer.com/10.1007/s00221-002-1247-x, doi:10.1007/s00221-002-1247-x [Article]

Biebl, B., Arcidiacono, E., Kacianka, S., Rieger, J. W., & Bengler, K. (2022). Opportunities and limitations of a gaze-contingent display to simulate visual field loss in driving simulator studies. Frontiers in Neuroergonomics, 3, 916169. https://www.frontiersin.org/articles/10.3389/fnrgo.2022.916169/full, doi:10.3389/fnrgo.2022.916169 [Article]

Biebl, B., & Bengler, K. (2021). I spy with my mental eye – analyzing compensatory scanning in drivers with homonymous visual field loss. In N. L. Black, W. P. Neumann, & I. Noy (Eds.), Proceedings of the 21st congress of the international ergonomics association (IEA 2021) (Vol. 221, pp. 552–559). Springer International Publishing. (Series Title: Lecture Notes in Networks and Systems)

Biebl, B., Kuhn, M., Stolle, F., Xu, J., Bengler, K., & Bowers, A. R. (2024). Knowing me, knowing you—a study on top-down requirements for compensatory scanning in drivers with homonymous visual field loss. PLOS ONE, 19(3), e0299129. https://dx.plos.org/10.1371/journal.pone.0299129, doi:10.1371/journal.pone.0299129 [Article]

Bosworth, R. G., & Dobkins, K. R. (1999). Left-hemisphere dominance for motion processing in deaf signers. Psychological Science, 10(3), 256–262. https://journals.sagepub.com/doi/10.1111/1467-9280.00146, doi:10.1111/1467-9280.00146 [Article]

Bowers, A. R. (2016). Driving with homonymous visual field loss: a review of the literature. Clinical and Experimental Optometry, 99(5), 402–418. https://www.tandfonline.com/doi/full/10.1111/cxo.12425, doi:10.1111/cxo.12425 [Article]

Bowers, A. R., Ananyev, E., Mandel, A. J., Goldstein, R. B., & Peli, E. (2014). Driving with hemianopia: IV. head scanning and detection at intersections in a simulator. Investigative Opthalmology & Visual Science, 55(3), 1540. http://iovs.arvojournals.org/article.aspx?doi=10.1167/iovs.13-12748, doi:10.1167/iovs.13-12748 [Article]

Bowers, A. R., Mandel, A. J., Goldstein, R. B., & Peli, E. (2010). Driving with hemianopia, II: Lane position and steering in a driving simulator. Investigative Opthalmology & Visual Science, 51(12), 6605. http://iovs.arvojournals.org/article.aspx?doi=10.1167/iovs.10-5310, doi:10.1167/iovs.10-5310 [Article]

Brookes, J., Warburton, M., Alghadier, M., Mon-Williams, M., & Mushtaq, F. (2020). Studying human behavior with virtual reality: The unity experiment framework. Behavior Research Methods, 52(2), 455–463. http://link.springer.com/10.3758/s13428-019-01242-0, doi:10.3758/s13428-019-01242-0 [Article]

Chen, X., McNamara, T. P., Kelly, J. W., & Wolbers, T. (2017). Cue combination in human spatial navigation. Cognitive Psychology, 95, 105–144. https://linkinghub.elsevier.com/retrieve/pii/S0010028516302043, doi:10.1016/j.cogpsych.2017.04.003 [Article]

Delpiano, R. (2021). Understanding the lateral dimension of traffic: Measuring and modeling lane discipline. Transportation Research Record: Journal of the Transportation Research Board, 2675(12), 1030–1042. https://journals.sagepub.com/doi/10.1177/03611981211031884, doi:10.1177/03611981211031884 [Article]

Donges, E. (1978). A two-level model of driver steering behavior. Human Factors: The Journal of the Human Factors and Ergonomics Society, 20(6), 691–707. https://journals.sagepub.com/doi/10.1177/001872087802000607, doi:10.1177/001872087802000607 [Article]

Ernst, M. O., & Banks, M. S. (2002). Humans integrate visual and haptic information in a statistically optimal fashion. Nature, 415(6870), 429–433.

Frissen, I., & Mars, F. (2014). The effect of visual degradation on anticipatory and compensatory steering control. Quarterly Journal of Experimental Psychology, 67(3), 499–507. https://journals.sagepub.com/doi/10.1080/17470218.2013.819518, doi:10.1080/17470218.2013.819518 [Article]

Ghosh, S. (2011). Typical coefficient of friction values for common materials. https://mechguru.com/machine-design/typical-coefficient-of-friction-values-for-common-materials/. (Accessed: January 16, 2025)

Giguere, A. P., Cavanaugh, M. R., Huxlin, K. R., Tadin, D., Fajen, B. R., & Diaz, G. J. (2025). The effect of unilateral cortical blindness on lane position and gaze behavior in a virtual reality steering task.

Giguere, A. P., Huxlin, K. R., Tadin, D., Fajen, B. R., & Diaz, G. J. (2024). Optic flow density modulates corner-cutting in a virtual steering task for younger and older adults. Scientific Reports, 14(1), 27693. https://www.nature.com/articles/s41598-024-78645-3, doi:10.1038/s41598-024-78645-3 [Article]

Horton, J. C., Economides, J. R., & Adams, D. L. (2021). The mechanism of macular sparing. Annual Review of Vision Science, 7(1), 155–179. https://www.annualreviews.org/doi/10.1146/annurev-vision-100119-125406, doi:10.1146/annurev-vision-100119-125406 [Article]

Iorizzo, D. B., Riley, M. E., Hayhoe, M., & Huxlin, K. R. (2011). Differential impact of partial cortical blindness on gaze strategies when sitting and walking – an immersive virtual reality study. Vision Research, 51(10), 1173–1184. https://linkinghub.elsevier.com/retrieve/pii/S0042698911001015, doi:10.1016/j.visres.2011.03.006 [Article]

Kasneci, E., Sippel, K., Aehling, K., Heister, M., Rosenstiel, W., Schiefer, U., et al. (2014). Driving with binocular visual field loss? a study on a supervised on-road parcours with simultaneous eye and head tracking. PLoS ONE, 9(2), e87470. https://dx.plos.org/10.1371/journal.pone.0087470, doi:10.1371/journal.pone.0087470 [Article]

Kim, H.-Y. (2015). Statistical notes for clinical researchers: post-hoc multiple comparisons. Restorative Dentistry & Endodontics, 40(2), 172. http://rde.ac/journal/view.php?doi=10.5395/rde.2015.40.2.172, doi:10.5395/rde.2015.40.2.172 [Article]

Kountouriotis, G. K., Floyd, R. C., Gardner, P. H., Merat, N., & Wilkie, R. M. (2012). The role of gaze and road edge information during high-speed locomotion. Journal of Experimental Psychology: Human Perception and Performance, 38(3), 687–702. http://doi.apa.org/getdoi.cfm?doi=10.1037/a0026123, doi:10.1037/a0026123 [Article]

Land, M., & Horwood, J. (1995). Which parts of the road guide steering? Nature, 377.

Landy, M. S., Maloney, L. T., Johnston, E. B., & Young, M. (1995). Measurement and modeling of depth cue combination: in defense of weak fusion. Vision research, 35(3), 389–412.

Lappi, O. (2014). Future path and tangent point models in the visual control of locomotion in curve driving. Journal of Vision, 14(12), 21–21. http://jov.arvojournals.org/Article.aspx?doi=10.1167/14.12.21, doi:10.1167/14.12.21 [Article]

Learmonth, G., Märker, G., McBride, N., Pellinen, P., & Harvey, M. (2018). Right-lateralised lane keeping in young and older british drivers. PLOS ONE, 13(9), e0203549. https://dx.plos.org/10.1371/journal.pone.0203549, doi:10.1371/journal.pone.0203549 [Article]

Logitech. (2025). G920 driving force racing wheel. Available from https://www.logitechg.com/en-us/products/driving/driving-force-racing-wheel.html

Mole, C. D., Kountouriotis, G., Billington, J., & Wilkie, R. M. (2016). Optic flow speed modulates guidance level control: New insights into two-level steering. Journal of Experimental Psychology: Human Perception and Performance, 42(11), 1818–1838. http://doi.apa.org/getdoi.cfm?doi=10.1037/xhp0000256, doi:10.1037/xhp0000256 [Article]

Newman, P. M., Qi, Y., Mou, W., & McNamara, T. P. (2023). Statistically optimal cue integration during human spatial navigation. Psychonomic Bulletin & Review, 30(5), 1621–1642. https://link.springer.com/10.3758/s13423-023-02254-w, doi:10.3758/s13423-023-02254-w [Article]

Papageorgiou, E., Hardiess, G., Mallot, H. A., & Schiefer, U. (2012). Gaze patterns predicting successful collision avoidance in patients with homonymous visual field defects. Vision Research, 65, 25–37. https://linkinghub.elsevier.com/retrieve/pii/S0042698912001757, doi:10.1016/j.visres.2012.06.004 [Article]

Parker, W. T., McGwin, G., Wood, J. M., Elgin, J., Vaphiades, M. S., Kline, L. B., et al. (2011). Self-reported driving difficulty by persons with hemianopia and quadrantanopia. Current Eye Research, 36(3), 270–277. http://www.tandfonline.com/doi/full/10.3109/02713683.2010.548893, doi:10.3109/02713683.2010.548893 [Article]

Patterson, G., Howard, C., Hepworth, L., & Rowe, F. (2019). The impact of visual field loss on driving skills: A systematic narrative review. British and Irish Orthoptic Journal, 15(1), 53. https://www.bioj-online.com/article/10.22599/bioj.129/, doi:10.22599/bioj.129 [Article]

Salvucci, D. D., & Gray, R. (2004). A two-point visual control model of steering. Perception, 33(10), 1233–1248. http://journals.sagepub.com/doi/10.1068/p5343, doi:10.1068/p5343 [Article]

Stein, N., Niehorster, D. C., Watson, T., Steinicke, F., Rifai, K., Wahl, S., et al. (2021). A comparison of eye tracking latencies among several commercial head-mounted displays. i-Perception, 12(1), 2041669520983338. https://journals.sagepub.com/doi/10.1177/2041669520983338, doi:10.1177/2041669520983338 [Article]

Szlyk, J. P., Brigell, M., & Seiple, W. (1993). Effects of age and hemianopic visual field loss on driving. American Academy of Optometry, 70(12), 1031–1037.

Tytarenko, M. (2023). Optimizing immersion: Analyzing graphics and performance considerations in unity3d VR development. Asian Journal of Research in Computer Science, 16(4), 104–114. https://journalajrcos.com/index.php/AJRCOS/article/view/374, doi:10.9734/ajrcos/2023/v16i4374 [Article]

VIVE. (n.d.). Vive pro specifications and user guide. Available from https://developer.vive.com/resources/hardware-guides/vive-pro-specs-user-guide/

Wood, J. M., Black, A. A., Mallon, K., Kwan, A. S., & Owsley, C. (2018). Effects of age-related macular degeneration on driving performance. Investigative Opthalmology & Visual Science, 59(1), 273. http://iovs.arvojournals.org/article.aspx?doi=10.1167/iovs.17-22751, doi:10.1167/iovs.17-22751 [Article]

